# Comparison of mechanical sorting and DNA metabarcoding for diet analysis with degraded wolf scats

**DOI:** 10.1101/2019.12.13.875898

**Authors:** Aimee Massey, Gretchen Roffler, Tessa Vermeul, Jennifer Allen, Taal Levi

## Abstract

DNA metabarcoding has become a powerful technique for identifying species and profiling biodiversity with the potential to improve efficiency, reveal rare prey species, and correct mistaken identification error in diet studies. However, the extent to which molecular approaches agree with traditional approaches is unknown for many species. Here, we compare diets from wolf scats profiled using both mechanical sorting and metabarcoding of amplified vertebrate DNA sequences. Our objectives were: (1) compare findings from mechanical sorting and metabarcoding as a method of diet profiling and (2) use results to better understand diets of wolves on Prince of Wales Island, a population of conservation concern. We predicted metabarcoding would reveal both higher diversity of prey and identify rare species that are overlooked with mechanical sorting. We also posited that the relative contribution of Sitka black-tailed deer (*Odocoileus hemionus sitkensis*) and beaver (*Castor canadensis*) would be overestimated with mechanical sorting methods because of the failure to account for the full diet diversity of these wolves. We found that there was substantial overlap in the diets revealed using both methods, indicating that deer, beaver, and black bear (*Ursus americanus*) were the primary prey species. However, there was a large discrepancy in the occurrence of beaver in scats (54% and 24% from mechanical sorting and metabarcoding, respectively) explained by the high rate of false positives with mechanical sorting methods. Metabarcoding revealed more diet diversity than mechanical sorting, thus supporting our initial predictions. Prince of Wales Island wolves appear to have a more diverse diet with greater occurrence of rare species than previously described including 14 prey species that contribute to wolf diet. Metabarcoding is an effective method for profiling carnivore diet and enhances our knowledge concerning the full diversity of wolf diets, even in the extremely wet conditions of southeast Alaska, which can lead to DNA degradation. Given the increasingly efficient and cost-effective nature of collecting eDNA, we recommend incorporating these molecular methods into field-based projects to further examine questions related to increased use of alternate prey coinciding with changes in abundance of primary prey and habitat alteration.

## Introduction

Animal scats are a vital tool for answering scientific questions related to animal behavior, diet, and species interactions. Traditionally, scat-based diet analysis has relied upon the mechanical processing and sorting of scat remains. This typically includes processing a scat to remove fecal material followed by meticulous sorting and identification of remaining hair and hard parts. However, diet analysis with mechanical sorting of scats has well-known biases (Lake et al. 2003); rare species or species that lack non-digestible hard parts are often overlooked or misidentified. In addition, some species are challenging to distinguish based on bone fragments or hair samples leading to additional misidentification. This is often the case for large mammals that consume prey tissue rather than whole individuals such that diagnostic hard parts like teeth and bones are frequently absent in scats. Metabarcoding of fecal DNA presents a new alternative method for diet analysis (Shehzad et al. 2012, De Barba et al. 2014, Kartzinel et al. 2015, McInnes et al. 2017, Eriksson et al. 2019). The DNA metabarcoding workflow includes extracting DNA from environmental samples, DNA amplification using ‘universal’ primers (Binladen et al. 2007), and mass-parallel sequencing of amplified product using next generation sequencing technologies. This process allows DNA barcodes from multiple species in a bulk sample to be sequenced simultaneously for an efficient and thorough profile of species present within an environmental sample (Valentini et al. 2009).

The utility of metabarcoding for informing important management objectives, where accurate taxonomic assignment and detection is paramount, is uncertain because unlike mechanical sorting (1) it is unknown how quality of inference from metabarcoding depends on acquiring relatively fresh scats with minimally degraded DNA, which can be challenging for rare taxa, and (2) it is not yet clear if the relative read abundance from metabarcoding can yield quantitative information that approximates the volume or biomass arising from each prey species. The degree to which relative read abundance (RRA) from DNA metabarcoding is correlated with the relative biomass of each prey species is a subject of substantial debate (Deagle et al. 2018, Pinol et al. 2018). Limited empirical research validating RRA against estimated biomass or volume from mechanical sorting informs this debate (Soininen et al. 2009, Thomas et al. 2017), although no studies have done so with terrestrial carnivores. Pinol et al. (2018) argued that metabarcoding results can only be interpreted quantitatively if amplification of DNA through PCR with universal primers is avoided because different amplification efficiencies among species can lead to poor representation of original biomass proportions. While this is often true for invertebrates, for which primer mismatch is common (Krehenwinkel et al. 2017), our 12S mtDNA primers rarely contains basepair mismatches for vertebrates and contain no mismatches for the taxa considered here (Appendix S1: Fig. S1). In addition, recent evidence suggests that as long as primer efficiency is high (no mismatches), the proportion of sequences arising from each species in metabarcoding (RRA) can produce semi-quantitative results (Kartzinel et al. 2015, Thomas et al. 2016, Krehenwinkel et al. 2017, Deagle et al. 2018). This could allow metabarcoding to approximate relative biomass or volume information similar to that produced by mechanical sorting of hard parts as well as frequency of occurrence (proportion of scats that contain each species). Nevertheless, the degree to which degraded scats yield suitable inference comparable to mechanical sorting is not currently well-understood because of a lack of formal comparisons between metabarcoding and mechanical sorting (Deagle et al. 2018, Pinol et al. 2018).

To provide this methods comparison, we focused on the Alexander Archipelago wolf (*Canis lupus ligoni*) as a case-study. The Alexander Archipelago wolf has been repeatedly petitioned for listing as threatened under the U.S. Endangered Species Act (ESA). These wolves occur in relative geographic isolation in southeast Alaska, where continued pressure from habitat loss, population decline of their primary prey, and wolf harvesting have raised concern about the future of the population. Wolf population estimates at regional scales in southeast Alaska have been based on expected Sitka black-tailed deer (*Odocoileus hemionus sitkensis*) abundance under the assumption that wolves are closely tied to the abundance of their primary prey. This is evident in the most recent ESA species status review where deer habitat quality metrics were used to project wolf abundance (U.S. Fish and Wildlife Service 2015).

The wolves on Prince of Wales Island (POW) (Fig. 1) were of particular concern in the most recent assessment (2015) because in addition to high levels of wolf harvest, POW has the highest rate of old-growth logging in southeast Alaska (Albert and Schoen 2013, Person and Brinkman 2017). Deer populations are predicted to decline as old-growth forests with palatable understory forbs and shrubs are converted into dense, even-aged, closed canopy forests (Alaback 1982, Schoen et al. 1988, Person et al. 1996, Farmer and Kirchhoff 2007, Gilbert et al. 2016, Person and Brinkman 2017, Porter 2018) that are strongly avoided by deer (Wallmo and Schoen 980, Kohira and Rexstad 1997, Gilbert et al. 2017). Deer were the most frequently occurring prey species for the Alexander Archipelago wolf based on previous research conducted on POW (Kohira 1995, Person et al. 1996, Kohira and Rexstad 1997). However, mechanical sorting of wolf scats has revealed other prey in significant quantities (Kohira 1995), and coastal wolves in this region can consume substantial quantities of salmon seasonally and other marine resources (Szepanski et al. 1999, Darimont et al. 2003, 2004, 2008a, Lafferty et al. 2014), suggesting that wolf population abundance may also be dictated by the availability of prey other than deer. Consequently, refining knowledge regarding the diet of wolves in the system has important implications for wolf management, potential ESA considerations, and forest management in southeast Alaska.

**Figure 1:**
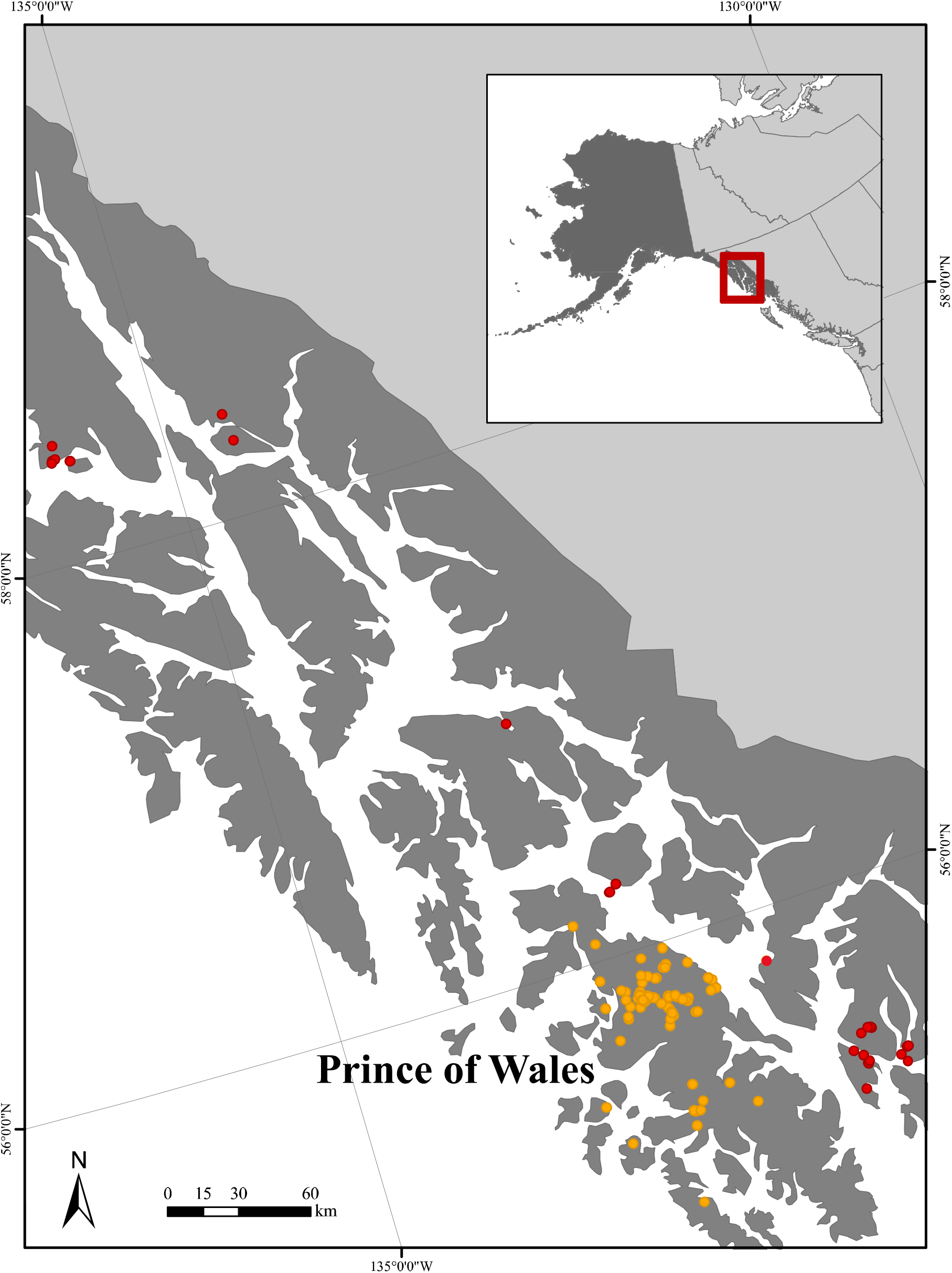
Study area map showing Alexander Archipelago in Southeast Alaska. Red and yellow points represent individual scat collection sites. Most scats collections were concentrated on Prince of Wales Island (yellow points).

Here we provide the first formal comparison of carnivore diet analysis from mechanical sorting and DNA metabarcoding using opportunistically collected scats across an assumed degradation spectrum in a temperate rainforest which is hostile to DNA preservation. We examined whether metabarcoding revealed a more diverse wolf diet than mechanical sorting, achieved increased taxonomic precision, and identified infrequently consumed prey species. We included both scats appearing highly degraded and those appearing fresh and assessed whether age of scats or biases introduced during the molecular processing affected the diet profile shown by metabarcoding. We additionally analyzed in detail Alexander Archipelago wolf diets with a particular focus on Prince of Wales Island to determine the prey profile of wolves and their dependence on deer.

## Materials and Methods

### Study area and field collection

Southeast Alaska lies within the Alexander Archipelago composed of over 2,000 named islands (Fig. 1) (Cook et al. 2006). This region receives between 130 – 400 cm of precipitation annually (Shanley et al. 2015) thus making it particularly inhospitable to the preservation of DNA in exposed environmental samples. The mainland is buttressed by the rugged Coast Mountains and extensive temperate rainforests at lower elevations. As a result of natural fragmentation and isolation, the North Pacific coast region supports many endemic plant and animal lineages, particularly on Prince of Wales Island, the largest island in the archipelago (Cook et al. 2006, MacDonald and Cook 2007, Smith 2016). Most of the forested area is within the Tongass National Forest managed by the U.S. Forest Service. This ecosystem hosts a diversity of mammals including iconic species such as Sitka black-tailed deer (*Odocoileus hemionus sitkensis*), American black bear (*Ursus americanus*), North American beaver (*Castor canadensis*), American marten (*Martes americanus*), mountain goat (*Oreamnos americanus*), Steller sea lion (*Eumetopias jubatus*), harbor seal (*Phoca vitulina*), and moose (*Alces alces*). Species distribution and assemblages vary among island and mainland areas of this region.

We collected wolf scats along wolf travel routes, near den sites, and on secondary roads during planned scat collection surveys during October 2014 - December 2015 (Fig. 1). We collected wolf scats primarily on Prince of Wales Island (55° 46’45.9480” N; 132° 49’ 4.7748” W) (n = 145), but also opportunistically collected samples in other mainland and island systems (n = 38). We estimated the age (fresh [<3 months] and old [>3 months]) of scat based on appearance, time since last site visit (Ciucci et al. 1996), and exposure time considering that scats decompose rapidly in rainforest environments (Wallmo et al. 1962, Ciucci et al. 1996, Darimont et al. 2008b) (Fig. 2). Collected wolf scats were stored in plastic bags, labeled with location, date, and perceived age of scat, and then frozen (−20°C). Frozen scats were shipped to Oregon State University for sample preparation and analysis.

**Figure 2:**
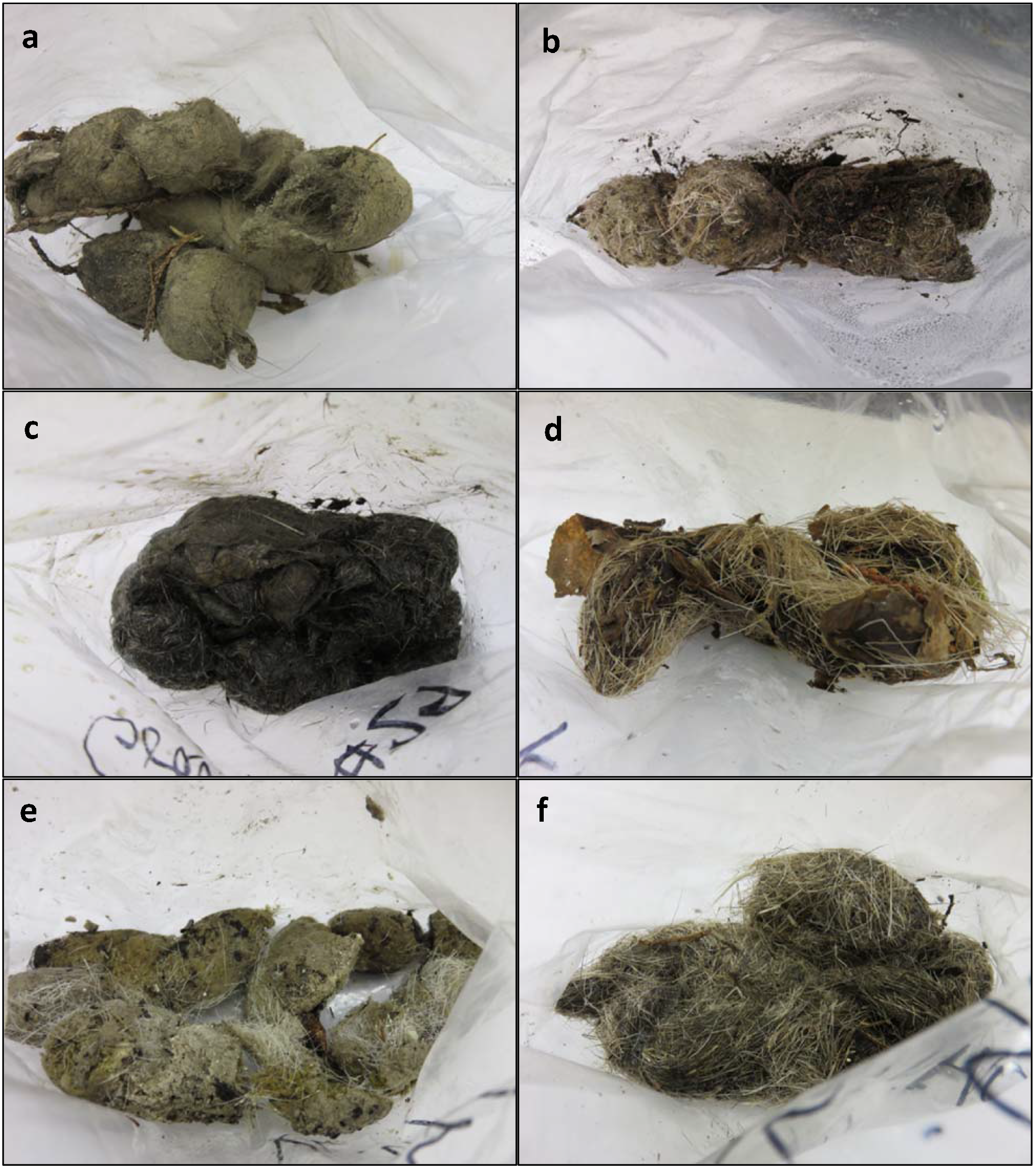
Examples of wolf scats collected in southeast Alaska near Prince of Wales Island. Left-sided panels (a, c, and e) are examples of fresh scats (< 3 months old) and the right-sided panels (b, d, f) are examples of old/degraded scats (> 3 months old). Age was determined by the collector; scats were collected throughout 2014 –2015.

### Mechanical sorting

We stored a subsample of each scat for later molecular analysis (sterilized forceps and razors were used to collect a sample from the middle section of each scat to minimize wolf DNA (Stenglein et al. 2010)), and then placed each scat in a mesh bag (1/8”) and soaked it in water for 48 hours in a mason jar. We power-washed the scat to remove as much remaining fecal matter as possible. The remaining contents (i.e., hair, bones, other hard parts) were put in a labeled paper bag and dried in an oven (at approximately 50°C) for at least five days. We weighed the processed scat material (hair, bone, scale, feather, etc.) and mechanically homogenized and sorted the remains by hand. On average, the fine-scale sorting took 3.6 hours per scat. We examined hairs under a microscope and compared to hair samples from the Alaska Fur ID project (Carrlee and Horelick 2011). We made slide mounts using clear nail polish to examine scale pattern and medulla diameter in order to identify species. Following identification, the slide was labeled with the species name and sample of origin. This exhaustive, fine-scale sorting (Fig. 3) ensured that even rare species could be identified. Along with species identification, we estimated the volume of each prey species as a proportion of estimated hard parts for a species in relation to all hard parts in an individual scat.

**Figure 3:**
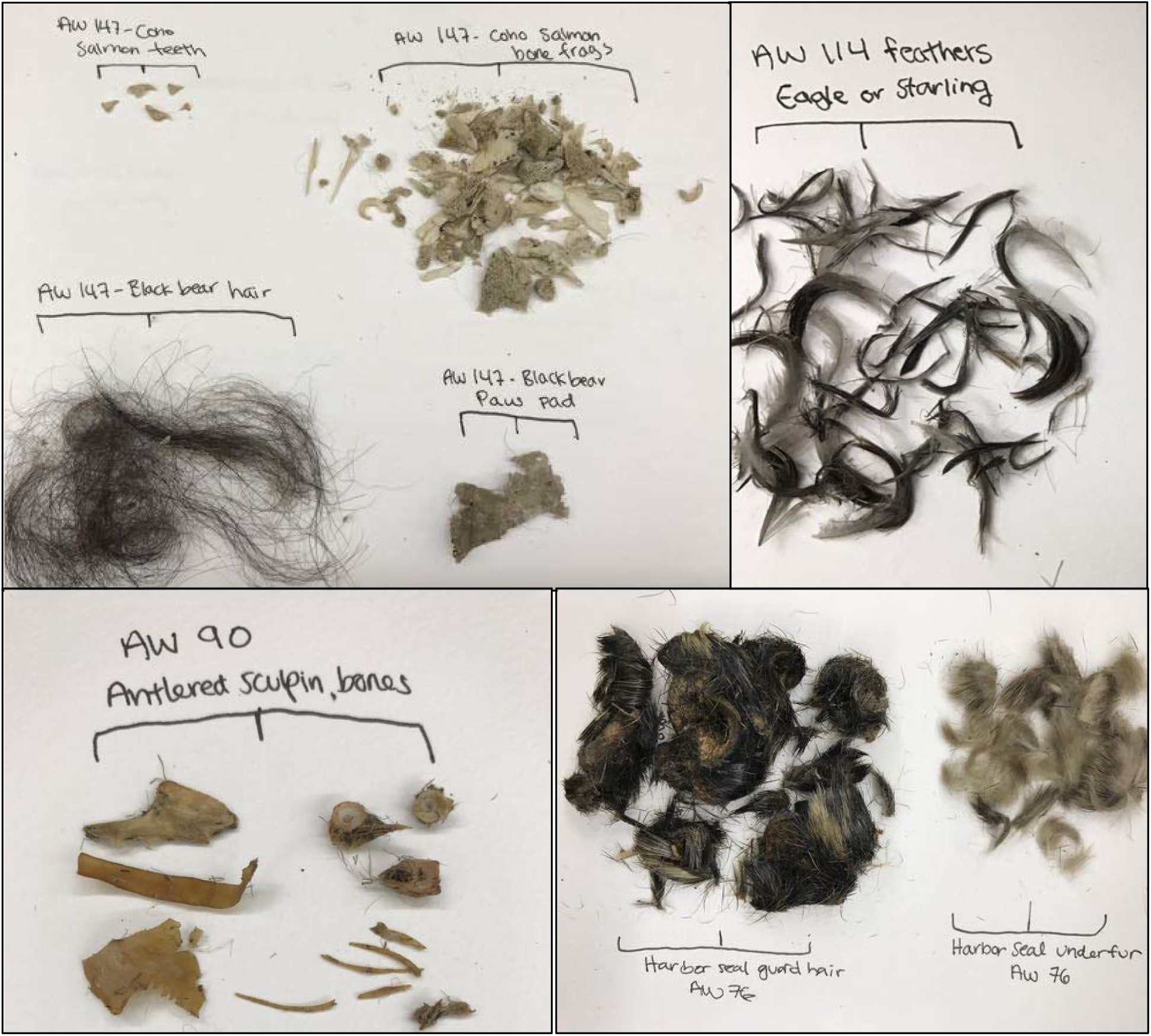
Photographs depicting examples of fine-scale mechanical sorting results of prey species in wolf scats collected in Southeast Alaska, 2014-2015. Starting from the top left panel and moving clockwise, species shown are salmon, black bear, bald eagle, harbor seal, and sculpin.

### Molecular analysis

Using the stored subsamples from each scat, we extracted DNA from each sample using the Qiagen DNeasy Blood and Tissue kit (Qiagen, Hilden, Germany) with slight modifications as follows: 500 ul Buffer ATL, 50 ul Proteinase K, and 1.0 mm Zirconia/Silica beads (BioSpec Products, Bartlesville, OK) were added to the 1.7 ml tube containing the scat. Samples were vortexed for 10 minutes at maximum speed prior to incubation at 56°C for 4-6 hours. The DNA was eluted in a total volume of 100 ul. A negative control was extracted with each round (approximately 17 samples) of DNA extraction to identify possible cross contamination.

Following DNA extraction, each sample was amplified in three separate reactions using the primer pair 12SV5F/12SV5R (Riaz et al. 2011). We used the forward primer (TTAGATACCCCACTATGC) as Riaz et al. (2011) but modified the first base pair of the reverse primer (YAGAACAGGCTCCTCTAG) to allow broader binding of vertebrate targets. These primers target approximately 100 base pairs in the 12S region of the vertebrate mitochondrial genome. The initial PCR was carried out using AmpliTaq Gold 360 Master Mix (Life Technologies, Carlsbad, CA). To label samples for multiplexing, we used 384 unique 8 bp dual matching indexes on the forward and reverse primers to eliminate contamination due to tag jumping by filtering reads that did not have identical indexes, and we included 3 bp of random nucleotides on the 5’end to increase sequence diversity and prevent degradation of indexes during subsequent blunt-ending and ligation steps. PCR reactions were carried out in a volume of 20 ul with 10 ul AmpliTaq Gold 360 Master Mix for a final concentration of 1x, 5 ul of forward and reverse primers for a final concentration of 0.25 uM, 3 ul of water, and 2 ul of DNA template. PCR cycling included initial denaturing at 95°C for 10 minutes, followed by 40 cycles of 95°C for 30 seconds, 58°C for 30 seconds, and 72°C for 30 seconds, with a final extension at 72°C for 7 minutes.

After the initial PCR, all PCR amplicons were cleaned using PCRClean DX solid-phase reversible immobilization magnetic beads (Aline Biosciences, Woburn, MA). Each PCR reaction was quantified using Accublue High Sensitivity dsDNA Quantitation kit (Biotium, Fremont, CA) and normalized to 6 ng/ul. Each group of 384 PCR products was then pooled into a single library, for a total of 3 libraries. Individual libraries were then tagged with an additional 6 base pair identifying index using the NEBnext Ultra II DNA Library Prep kit (New England Biolabs, Ipswich, MA). Pooled samples were analyzed on a Bioanalyzer to confirm fragment size. The libraries were then sequenced on one lane of Illumina HiSeq 3000 2 × 150 bp PE at the Center for Genome Research and Biocomputing at Oregon State University.

### Sequence analysis

Raw sequence reads were analyzed using a bioinformatics pipeline designed to trim and sort the sequence reads according to scat sample identification. An outline of the bioinformatic process is as follows: (1) raw reads were paired using PEAR (Zhang et al. 2013); (2) followed by demultiplexing using 8 basepair index sequences unique to each sample (mismatches discarded); (3) lastly, sequences from each sample were clustered by 100% similarity and taxonomically assigned using BLAST against 12S vertebrate sequences in GenBank and from a custom 12S database.

Similar to the step-wise methods used by De Barba et al. (2014), a series of filtering and quality control measures were carried out on taxonomically assigned sequences. We initially removed sequences that were identified to be *Canis* spp. and contaminants based on read counts in no-template controls (which contained primarily human contamination). We then removed sample replicates that failed to amplify during PCR which included sample replicates with fewer than a total of 400 sequence reads. We compared taxonomic assignments with known fauna of southeast Alaska (MacDonald and Cook 2007) to replace non-regional species identified with BLAST with closely-related regional taxa. We then excluded prey items occurring in fewer than 2 of 3 PCR replicates. Finally, we combined those sample replicates that amplified so that sequence reads were totaled for each species within a sample and over the entire sample and eliminated sequences that appeared in less than 1% of the total reads for an individual sample.

### Age of scats

Prior to processing, we observed marked differences between the appearance and quality of scats (Fig. 2). We performed t-tests to determine whether the perceived age of a scat made during field collection correlated with either the average quantity of DNA (ng/ul) in a sample (measured post normalization using Accublue High Sensitivity dsDNA Quantitation kit (Biotium, Fremont, CA)), the total number of sequence reads in a sample including the wolf defecator, or the total number of sequence reads excluding wolf.

### Frequency of occurrence

We used both frequency of occurrence (FOO) and metrics of relative abundance (see below) to describe the occurrence of prey in wolf diet. FOO was calculated to determine which prey species were present and how often they were present based on the number of samples. For mechanical sorting methods, a species was present if there was evidence (including trace elements) of a prey species (e.g., hair, bone, scales, etc.) within a scat sample. FOO was then calculated as the proportion of scats in which a prey species occurred. For metabarcoding, a species occurrence was determined by whether sequence reads for a particular species were found in an individual scat after quality control measures. We compared FOO from mechanical sorting and metabarcoding using the subset of scats analyzed by both methods (n = 104), but we additionally present diet analysis from all scats collected on Princes of Wales Island and close surrounding islands (n = 118 metabarcoding; n = 98 mechanical sorting) to describe diet on POW.

To analyze discrepancies between species present in samples with mechanical sorting and not found with metabarcoding, we used generalized logistic regression with logit link to explore whether false positives from mechanical sorting or false negatives generated from metabarcoding best explained the absence of species. Statistical analyses were conducted in the R statistical program using the ‘stats’ package (R Core Team 2018). We reasoned that false negatives could arise if scats contained poor quality DNA or sequencing depth was insufficient. We therefore fit three separate logistic regression models using average DNA quantity per sample (across the three replicates PCRs), total number of sequence reads prior to quality control and including wolf sequence reads, and total number of sequences reads post quality control and not including wolf sequences reads as univariate predictors in each model. In our analysis, zeroes were defined as an absence in metabarcoding where mechanical sorting had indicated an occurrence of a particular species in a sample; one indicated where metabarcoding was in agreement with occurrence found in mechanical sorting. Therefore, positive coefficients imply an increasing rate of proper assignment as DNA quality or sequencing depth increases. The absence of such an effect would suggest that mismatch between metabarcoding and mechanical sorting is unlikely to be due to false negatives by metabarcoding.

### Relative abundance

To test whether metabarcoding and mechanical sorting yield similar metrics for relative abundance of a prey species within a scat, we compared percent estimated volume from mechanical sorting with the relative read abundance (RRA) from metabarcoding. RRA for each species *i* was calculated as

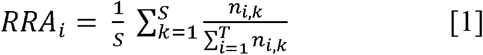

where *n*_*i,k*_ is the number of sequences of prey species *i* in sample *k*, *S* is the total number of samples, and *T* is the total number of species. We compared estimated volume of a prey species from mechanical sorting with RRA from metabarcoding using simple linear regression (R Core Team 2018).

For both the frequency of occurrence and relative abundance analyses we additionally revisited results from scats with mismatches from metabarcoding and mechanical sorting to assess whether metabarcoding found many sequence reads of an alternative species that was incorrectly assigned by mechanical sorting and was thus likely a false positive.

## Results

### Age of scats

Purportedly fresh scats contained significantly more total sequence reads on average (μ_fresh_ = 269,514 ± 173,902) compared to the total number of reads from degraded wolf scats (μ_degraded_ = 200,378 ± 135,646) (t = 2.09, df = 85, p-value = 0.039). Likewise, fresh scats (μ_fresh_ = 139,939 ± 135,858) had significantly more wolf sequence reads than degraded scats (μ_degraded_ = 52,411 ± 75,531) (t = 3.80, df = 73.73, p-value < 0.001). However, we found no significant difference between degraded and fresh scats when considering only reads from prey items (excluding any wolf DNA reads), although degraded scats yielded a greater average number of non-wolf reads per sample than fresh scats (μ_degraded_ = 147,966 ± 125,223; μ_fresh_ = 129,575 ± 124,124; t = −0.69, df = 84.18, p-value = 0.49) (Fig. 4). Fresh scats had a higher average DNA quantity post PCR (ng/ul; μ_degraded_ = 4.12 ± 1.97; μ_fresh_ = 4.55 ± 2.20) but the difference was not statistically significant (t = 0.97, df = 85.93, p-value = 0.33).

**Figure 4:**
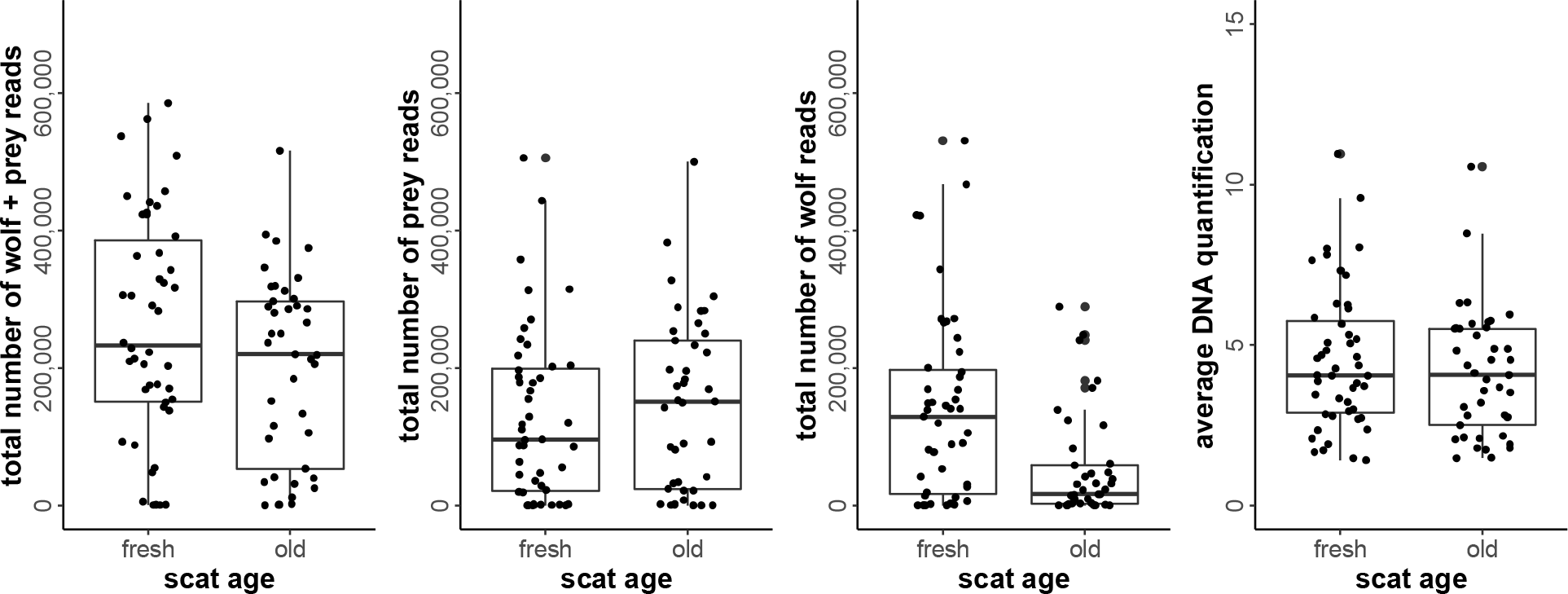
Boxplots depicting the total number of reads and the DNA quantity (measured post normalization) for scat samples binned by the age of the scat.

### Comparing wolf diet by mechanical sorting and metabarcoding – frequency of occurrence

We compared wolf diet from 104 scat samples that were analyzed with both mechanical sorting and metabarcoding. Metabarcoding revealed a number of rare species that were not found using mechanical sorting methods and thus revealed greater dietary diversity (Fig. 5). Species that were found with metabarcoding methods but were absent when using mechanical sorting methods include: duck (*Anas* spp.), dusky grouse (*Bonasa umbellus*), elk (*Cervus elaphus*), raven (*Corvus* species), Northern collared lemming (*Dicrostonyx groenlandicus*), Steller sea lion (*Eumetopias jubatus*), American marten (*Martes americana*), and American red squirrel (*Tamiasciurus hudsonicus*). Mechanical methods identified moose (*Alces alces*) in a single scat where metabarcoding did not, although moose was identified by metabarcoding in this particular scat prior to quality filtering.

**Figure 5:**
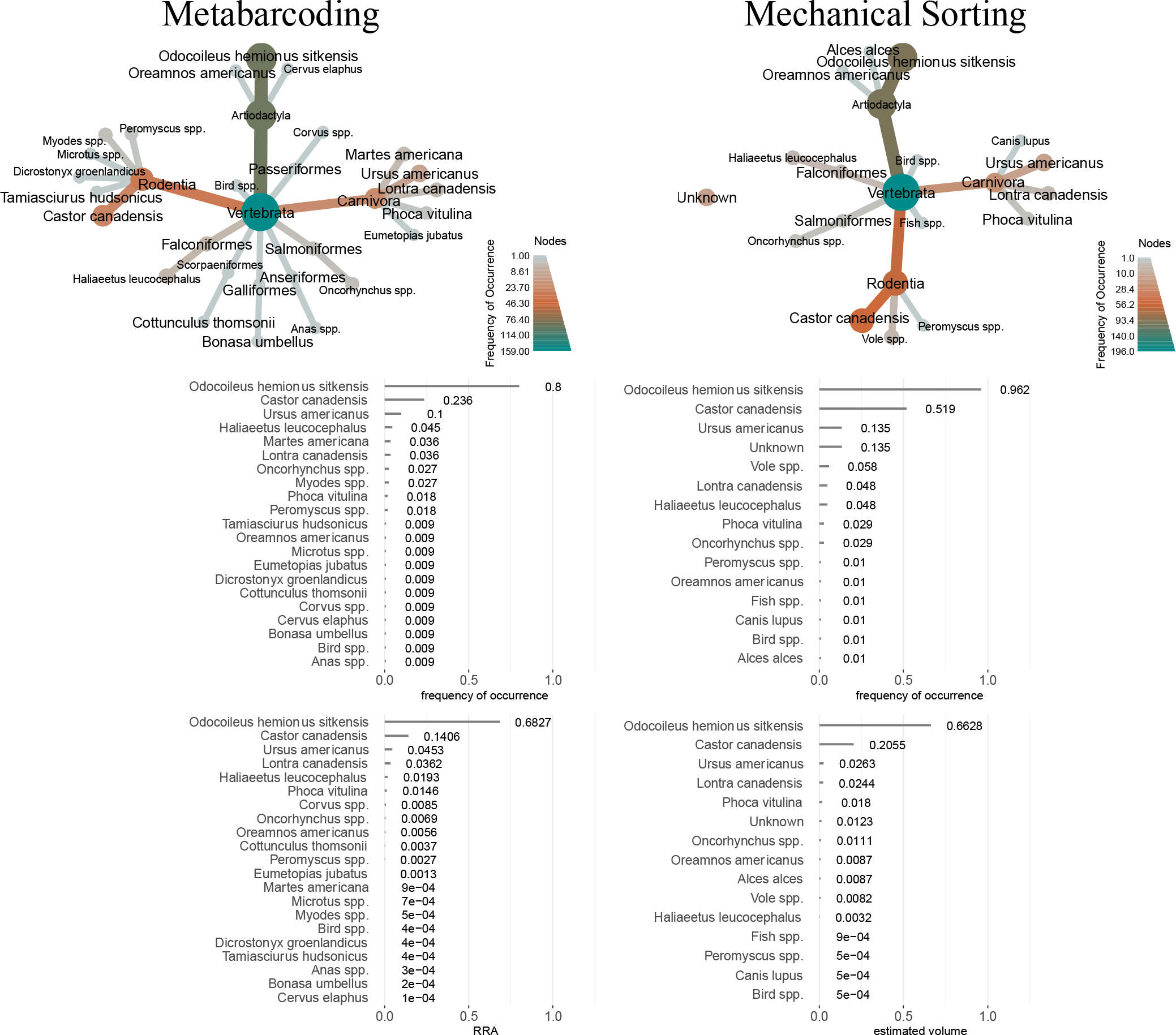
Diet summary from analysis of wolf scats (Southeast Alaska, 2014-2015) using (a) metabarcoding methods and (b) mechanical sorting methods. For the diet trees, each branch and terminal node represent a prey species identified in the wolf scats with the size and color of the branch showing the number of occurrences of that prey species. Frequency of Occurrence and RRA and estimated volume are compared.

Frequency of occurrence (FOO) (Fig. 5) results were similar with both methods. However, there was substantial discrepancy between the primary prey species (Sitka black-tailed deer) and the secondary prey species (beaver) between metabarcoding and mechanical occurrence results. The occurrence of deer was greater in the mechanical sorting results (FOO_mech_ = 0.962) compared to metabarcoding results (FOO_MB_ = 0.8) and the occurrence of beaver was twice as frequent in the mechanical sorting (FOO_mech_ = 0.519) results compared to metabarcoding (FOO_MB_ = 0.236).

Logistic regression to assess mismatch between metabarcoding and mechanical sorting revealed that neither average DNA quantity, total sequence reads, nor total sequence reads of prey (i.e. excluding wolf) were associated with failing to detect species that were identified by metabarcoding (Table 1). However, contrary to predictions, increasing number of prey sequence reads (i.e. excluding wolves) was associated with increasing mismatch with beaver occurrences detected by mechanical sorting (p = 0.025), which suggests that the error was due to misassignment by mechanical sorting rather than by metabarcoding. Thirty-two of the 59 beaver occurrences had disagreement between mechanical sorting and metabarcoding results. Notes and hair slides taken during mechanical sorting showed that 18 of the 32 mismatches could be attributed to false positives generated from mechanical sorting. In addition, a substantial number of definitive deer occurrences (i.e. high relative read abundance for deer) were mistakenly assigned to beaver by mechanical sorting (Fig. 6), further suggesting that mismatch between methods was due to misassignment by mechanical sorting.

**Table 1:** Summary statistics for all generalized logistic regression models. Predictor variables include average DNA quantity per sample (avg DNA quant), total number of sequence reads prior to quality control and including wolf sequence reads (reads with wolf), and total number of sequences reads post quality control and not including wolf sequences reads (reads no wolf). Models were tested against all mechanically sorted samples that had a positive occurrence for a species and against all mechanically sorted samples that had a positive occurrence for beaver.

| Model | Estimate | SE | z value | Pr(>\|z\|) |
| --- | --- | --- | --- | --- |
| spp.presence ~ avg DNA quantity | 0.13 | 0.14 | 0.966 | 0.33 |
| spp.presence ~ total reads with wolf | 4.16e-07 | 1.5e-06 | 0.273 | 0.79 |
| spp.presence ~ total reads no wolf | -1.5e-07 | 1.63e-06 | -0.095 | 0.93 |
| beaver.presence ~ avg DNA quantity | -0.034 | 0.139 | -0.25 | 0.81 |
| beaver.presence ~ total reads with wolf | 4.4e-07 | 1.78e-06 | 0.25 | 0.81 |
| beaver.presence ~ total reads no wolf | -5.85e-06 | 2.61e-06 | -2.24 | 0.025* |

**Figure 6:**
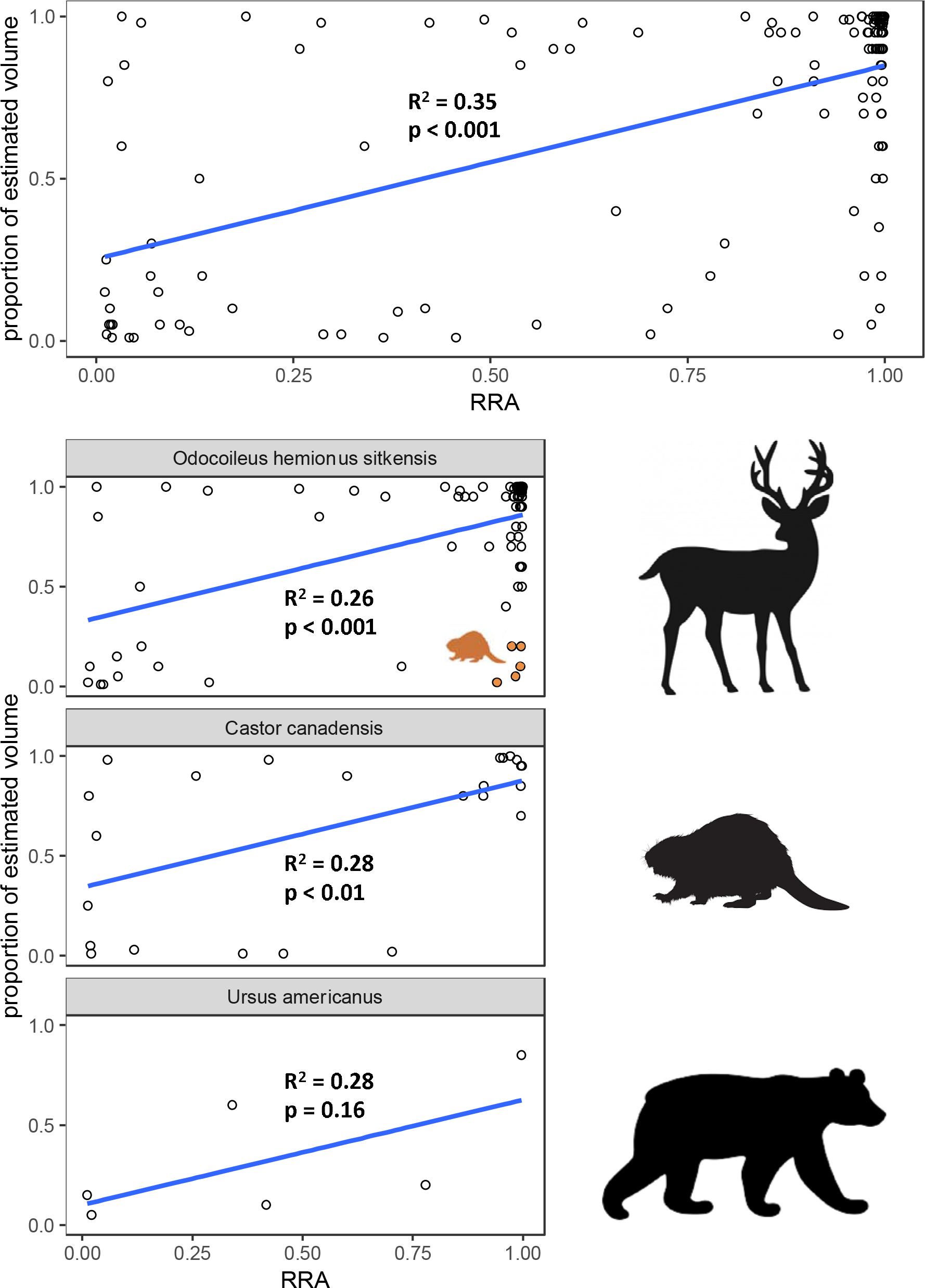
Correlation between relative read abundance data for metabarcoding methods and estimated volume for mechanical sorting methods by scat sample for the three most prevalent prey species from wolf scats, Southeast Alaska, 2014-2015. Estimated volume is measured as the proportion of a prey species consumed per scat relative to the whole scat. RRA is the relative read abundance. Data points highlighted in brown show samples where deer was thought to be mistakenly identified as beaver in mechanical sorting.

### Comparing wolf diet by mechanical sorting and metabarcoding – relative read abundance

There was minimal discrepancy between RRA of primary prey species (metabarcoding) and their estimated volume in scats (mechanical sorting); the difference between RRA and estimated volume for deer was 2% (RRA_deer_ = 68.3%; estimated volume_deer_ = 66.3%) and for beaver it was less than 7% (RRA_beaver_ = 14.1%; estimated volume_beaver_ = 20.5%). For the rarer species, we found a close association (within 2%) between the RRA and the estimated volume for that species.

The estimated volume from mechanical sorting was positively correlated with RRA of deer (*β* = 0.53; R^2^ = 0.26; *p* < 0.01, n = 87), beaver (*β*= 0.57; R^2^ = 0.28; *p* < 0.01, n = 25), and black bear (*β* = 0.80; R^2^ = 0.28; *p* = 0.17, n = 6) (Fig. 6), supporting a positive but variable relationship between the volume of parts of a particular species found in the physical scat and the proportion of DNA sequence reads for that species. However, substantial variability is likely due to species misidentification by mechanical sorting such as deer falsely identified as beaver (Fig. 6).

### Prince of Wales

Metabarcoding of scats found only within Prince of Wales Island (POW) (Fig. 7) revealed 14 species (Supplementary table) including Sitka black-tailed deer (FOO_MB_POW_ = 0.852), beaver (FOO_MB_POW_ = 0.231), and black bear (FOO_MB_POW_ = 0.157) were the most common prey items (Fig. 7). Other common prey species were salmon (*Oncorhynchus* spp.) (FOO_MB_POW_ = 0.056), American marten (*Martes americana*) (FOO_MB_POW_ = 0.046), North American river otter (*Lontra canadensis*) (FOO_MB_POW_ = 0.037), and bald eagle (*Haliaeetus leucocephalus*) (FOO_MB_POW_ = 0.019). Additional prey items in less than 1% of scats include American red squirrel (*Tamiasciurus hudsonicus*), deermouse (*Peromyscus* spp.), vole (*Myodes* and *Microtus* spp.), dusky grouse (*Bonasa umbellus*), duck (*Anas* spp.), and unidentified bird species.

**Figure 7:**
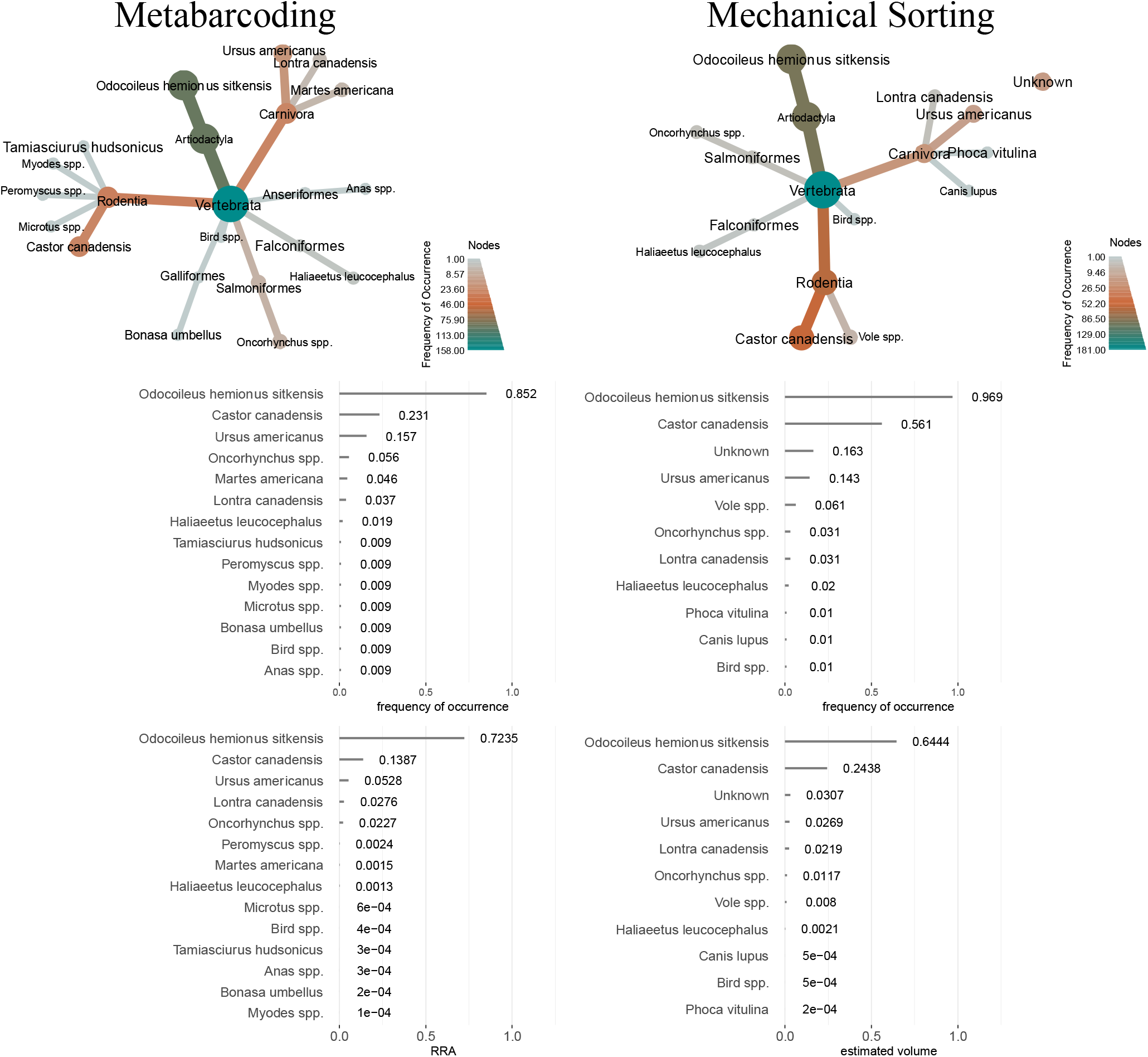
Diet diversity, frequency of occurrence (FOO), and RRA and estimated volume found with a) metabarcoding results and b) mechanical results for scats found on Prince of Wales Island, Alaska.

Mechanical sorting revealed a total of 10 prey species (Fig. 7), including harbor seal which was not found with metabarcoding for the POW samples. However, it should be noted that for this sample, mechanical sorting estimated only 2% harbor seal and metabarcoding instead found otter, which could have been mistaken for harbor seal during sorting. Deer (FOO_mech_POW_= 0.969) and beaver (FOO_mech_POW_ = 0.561) (the two primary prey species) showed greater FOO compared to metabarcoding, although mechanical sorting did not show any American marten and had a lower FOO of salmon species compared to the metabarcoding results. There was also substantial occurrence of material from unknown species in the mechanical results (FOO_mech_POW_ = 0.163) that is not seen with metabarcoding.

## Discussion

DNA metabarcoding has emerged as a novel method for diet analysis because of the ability to reveal rare or difficult to identify species (Shehzad et al. 2012, De Barba et al. 2014, Berry et al. 2015, Srivathsan et al. 2015, Kartzinel et al. 2015, McInnes et al. 2017, Buglione et al. 2018). However, substantial uncertainty remains as to whether inference from mechanical sorting and DNA metabarcoding produce comparable results, particularly if scats are of uncertain age and quality. Our results suggest that excluding purportedly degraded scats from DNA metabarcoding analyses does not improve inference about diet. Perceived fresh scats contained on average a greater number of reads per scat when including wolf sequence reads, but there was no significant difference in the average number of reads between fresh and degraded scats when only including reads from prey species (Fig. 4). The average quantity of DNA was also not significantly different between fresh and degraded scats; this is likely because fresh scats contained more fecal material relative to hair and bone, and total DNA quantity per sample is normalized prior to sequencing such that abundant wolf DNA leads to dilution of prey DNA. Many degraded scats were primarily clusters of hair and bone that were washed of fecal material. Importantly, these results suggest that metabarcoding is sensitive enough to determine prey assemblages in degraded scats and thus scat collection and processing should not be predicated upon perceived scat quality.

FOO and RRA metrics were qualitatively similar among methods. RRA of each species was significantly correlated with estimated volume determined with mechanical sorting (p_all_ < 0.01, p_deer_ < 0.01, p_beaver_ < 0.01, p_bear_ = 0.16) suggesting that RRA can be a reasonable proxy for volume of prey species obtained from mechanical sorting (which in turn could be used to estimate relative biomass using biomass equations that correct for body size (Weaver 1993)). Both mechanical sorting and metabarcoding agreed that Sitka black-tailed deer was the primary prey item, followed by beaver, and then black bear as suggested by previous research in this region (Kohira, 1995; Kohira & Rexstad, 1997) (Fig. 5). However, both deer and beaver occurred substantially more frequently in mechanically sorted scats than in metabarcoded scats. The divergence between the two methods examined in our study was more substantial for beaver which were identified mechanically in 52% of scats while only detected by metabarcoding in 24%.

We closely examined scats that were mismatched (i.e. the prey species was found in a scat during mechanical sorting but not found in the same scat with DNA metabarcoding) with a focus on beaver to assess whether mismatches were due to false positives produced from mechanical sorting or false negatives produced from metabarcoding. Eighteen of the 32 mismatched samples show evidence of false positive resulting from mechanical sorting. In these scats, beaver was thought to be present, but notes during sorting specified uncertainty that these small amounts of unknown hair samples could also be attributed to deer or black bear. In fact, we found that in all mismatched beaver samples metabarcoding showed a high RRA of deer and mechanical sorting found low volume of deer, strongly suggesting that mechanical sorting mis-assigned deer hair to beaver as the primary prey species in that scat (highlighted in Fig. 6). Our logistic regression analysis additionally suggests that these errors resulted from mis-assignment by mechanical sorting rather than metabarcoding (Table 1).

Why do we see these potential false positives generated from mechanical sorting? One explanation is that relying on mechanical sorting of scats results in the overestimation of primary prey species (i.e. deer and beaver) due to search image bias. Mechanical sorting can lead to mislabeling difficult to identify parts as common species rather than an infrequently detected species because the researcher is accustomed to encountering the common prey species. The pronounced difference seen in beaver FOO could also be attributed to the difficulty in distinguishing between beaver and guard hair from other species such as deer and black bear (Fig. 8).

**Figure 8:**
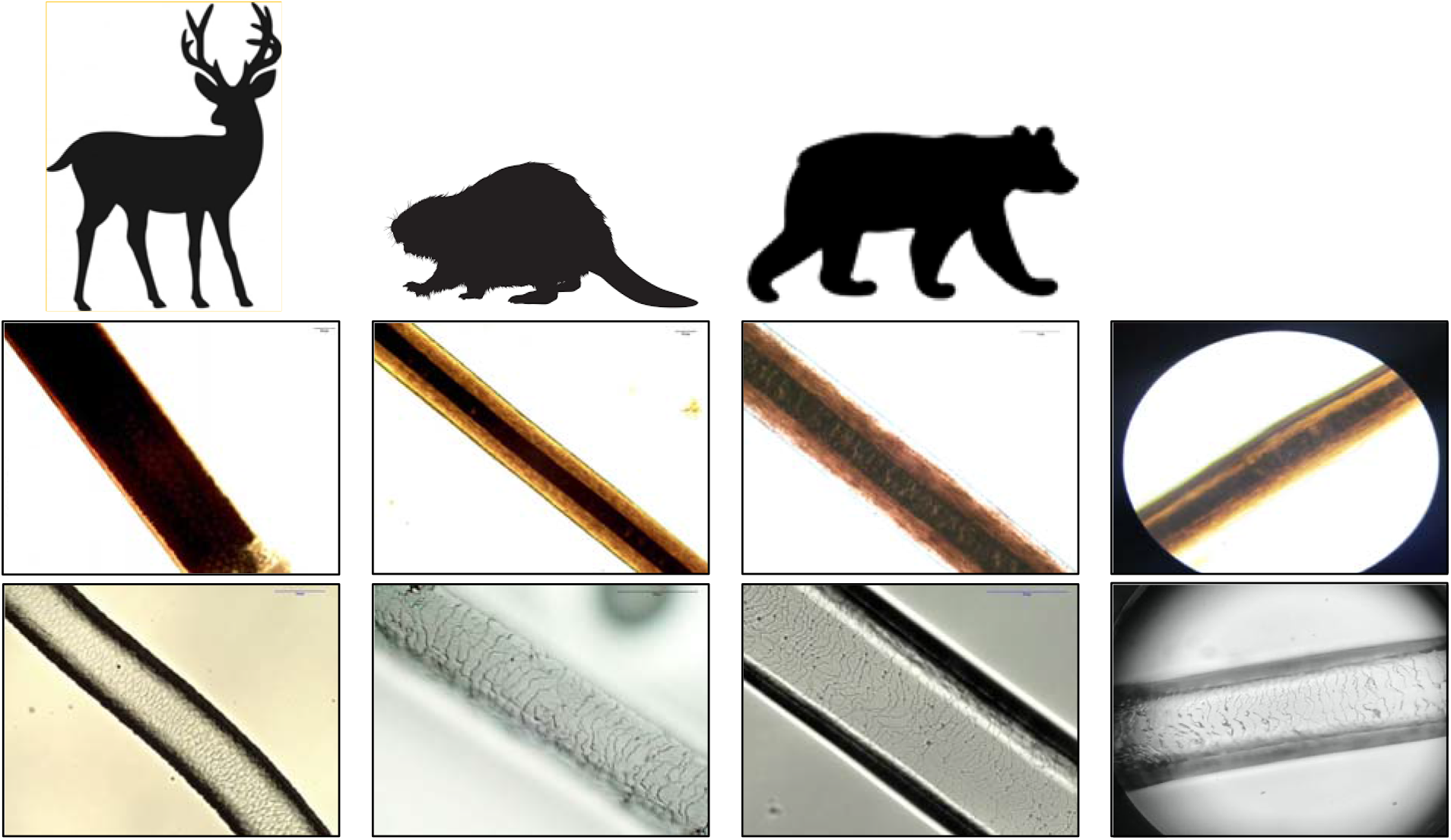
Panel of hair samples. The top row shows examples of guard hairs from the Alaska Fur ID project of Sitka black-tailed deer, beaver, and black bear. The bottom row shows examples of scale pattern from scale casts from the Alaska Fur ID project of Sitka black-tailed deer, beaver, and black bear (left to right). The last panel in each row is an example of a difficult to identify hair and scale pattern from a wolf scat sample, Southeast Alaska 2014-2015.

The remaining 14 of the 32 beaver mismatches were attributed to false negatives generated by metabarcoding; we concluded this because beaver was verified to have occurred in mechanical sorting but was absent from metabarcoding results. However, for 10 of these scats beaver occurred in the metabarcoding results prior to quality filtering that removed prey that occurred in fewer than 2 of 3 PCR replicates and with fewer that 1% of the total reads (importantly, beaver was nearly absent from our negative controls), which had the effect of underestimating prey items that occurred in only a small portion of a scat. It is important to note that our conservative quality filtering thresholds following De Barba et al. (2014) led to some of these false negatives at the expense of false positives. Thus, it is imperative to explicitly reason through quality control protocols to balance false positives and false negatives when using bioinformatically-generated metabarcoding data.

There was also divergence in the detection of rare species among methods. Although metabarcoding revealed several clear false negatives, this was substantially more common with mechanical sorting where 8 species in final metabarcoding results were not found by mechanical sorting for the same subset of samples. In particular, American marten, Northern collared lemming, and a number of bird species were missing from mechanical sorting but evident in the metabarcoding results. This conclusion supports our initial prediction that metabarcoding would be more advantageous in identifying rare species.

### POW wolf diet – policy and management

The issue of what wolves eat and how much is an important question in southeast Alaska and in particular on Prince of Wales Island where there are concerns about the long-term viability of wolves given the trophic linkage between wolves, Sitka black-tailed deer, and old-growth forest. The population of Sitka black-tailed deer is expected to decline in this region with continued logging of old-growth forests (Person and Brinkman 2017). Given this, wolf populations are predicted to decline and these declines are most significant under scenarios where wolves rely heavily on deer in the future (Gilbert et al. 2016).

Our study shows the promise of eDNA and metabarcoding methods to examine wolf diet diversity and diet changes. Comparing our results with previous work indicated that the occurrence of the primary prey species (Sitka black-tailed deer) is comparable on POW; Person et al. (1996) reported a >90% occurrence while we report 85.2% occurrence using DNA metabarcoding and 96.9% occurrence using mechanical sorting. However, the occurrence of beaver is greater compared to previous work; the frequency of occurrence of beaver was 13.7% (Person et al. 1996) and 31% (Kohira and Rexstad 1997), whereas we report 23.1% occurrence using DNA metabarcoding and 56.1% using mechanical sorting. These previous studies found that aside from Sitka black-tailed deer, beaver, and black bear, the only significant other prey were small mustelid species, river otter, and fish. Our results show a diverse diet with 14 total prey species identified from mechanical sorting that contribute to wolf diet on POW (Fig. 7), which more closely resembles the diversity found by Darimont et al. (2004) in their study of wolf diet using scats along the coastal region in British Columbia. Importantly, our findings suggest that metabarcoding was able to reveal the breadth of Alexander Archipelago wolf diet diversity more accurately than mechanical sorting. (24 vs. 14 refer to Appendix S1: Table S1).

Continued diet analysis using metabarcoding of wolf scats found on POW could reveal whether this increase in diversity is due to the increased power in the method used (metabarcoding vs. mechanical sorting), or if wolves are beginning to exhibit increased opportunistic predation on species other than Sitka black-tailed deer. Given that we also found greater dietary diversity using mechanical sorting compared to results using the same methods from the mid-1990’s points towards a potential dietary shift in wolves on POW (Kohira and Rexstad 1997). The rate of clear-cut logging in this region peaked during the late 1980’s and 1990’s and while this rate has slowed in recent years, a total of nearly 30% of old-growth forests have been logged on POW (U.S. Fish and Wildlife Service 2015). Because young-growth stands older than 25 years are the least productive in terms of deer forage (U.S. Fish and Wildlife Service 2015), the effects of potential deer abundance decline on wolf populations are only just being realized. As well-known diet generalists, it remains to be seen whether wolves on POW are resilient to landscape-level ecological changes expected from old-growth logging.

Metabarcoding has revealed a more diverse and precise diet for wolves on POW and in southeast Alaska, potentially pointing towards these wolves making greater use of alternate prey. In general, DNA metabarcoding can be used as a tool to reliably describe diet for other carnivore species. Even in a hostile environment for the preservation of eDNA, we have shown that DNA metabarcoding is an effective and powerful method for describing carnivore diet. Diet analysis remains one of the most important avenues of wildlife study as it is a necessary component of understanding species interactions, predator-prey dynamics, and the biodiversity of systems. This nuanced profiling of diet is especially important as vulnerable wildlife populations face continued habitat loss and degradation, and thus we can use changes in diet can as potential indicators of environmental health.

## Acknowledgements

This research was funded by the Alaska Department of Fish and Game and by Oregon State University. The authors would like to thank all those involved with collecting wolf scats including T. Bentz, S. Bethune, D. Gregovich, J. Jemmison, M. Kampnich, K. Larson, J. Manuel, G. Roffler, J. Reeves, S. Sell, Y. Shakeri, and R. Slayton as well as C. Cousins, D. Martinez, and A. Pepper for the assistance with the mechanical sorting of scats. The Center for Genome Research and Biocomputing (CGRB) at Oregon State University provided both access to the Illumina HiSeq 3000 for all metabarcoding and support for initial bioinformatic analysis.

