## Supplemental for "Comparison of mechanical sorting and DNA metabarcoding for diet analysis with degraded wolf scats"

**Appendix S1**

**Text S1**

*Full Wolf Diet Diversity*

A subset (n = 129) of the 183 total scats were processed and sorted using mechanical methods. Mechanical sorting revealed 14 known prey species (Table S1). Three % of the sorted scat material volume and 10% of total occurrences was labeled unknown or undetermined. We found evidence of wolf hard parts in one scat sample suggesting part of a wolf was digested by the wolf defecator, which could result from a wolf killing another wolf during territorial defense (Marhenke III 1971, Cassidy et al. 2015). Mechanical sorting revealed evidence of several uncommon species in the coastal wolf diet, including hard parts from harbor seal (n = 3), salmon (n = 3), sculpin (n = 1), bald eagle (n = 5), and two taxa of small mammals including a *Peromyscus* species (n = 1) and a vole species (n = 7) (Figure 3).

All 183 scats were analyzed using molecular methods. The number of paired sequence reads was 44,034,457 for the entire sample dataset. After quality control steps, the final dataset used for diet analysis had 26,199,194 sequence reads from 138 samples. 24 wolf prey species were identified from metabarcoding methods (Table S1).

**Table S1:** Total prey species diversity measured from presence/absence in 129 mechanically sorted scats and 138 molecularly processed scats from Alexander Archipelago wolves in Southeast Alaska, 2014-2015.

| Species | Mechanical | Metabarcoding |
| --- | --- | --- |
| *Alces alces* | y | y |
| *Anas* spp. |  | y |
| Bird spp. | y | y |
| *Bonasa umbellus* |  | y |
| *Canis lupus* | y | unknown |
| *Castor canadensis* | y | y |
| *Cervus elaphus* |  | y |
| *Corvus* spp. |  | y |
| *Cottunculus thomsonii* | y* | y |
| **Labeled as fish spp. for mech* |  |  |
| *Dicrostonyx groenlandicus* |  | y |
| *Eumetopias jubatus* |  | y |
| *Erethizon dorsatum* |  | y |
| *Gallus gallus* |  | y |
| *Haliaeetus leucocephalus* | y | y |
| *Lontra canadensis* | y | y |
| *Martes americana* |  | y |
| *Microtus* spp. | y* | y |
| *Myodes* spp. |  | y |
| **Labeled as vole spp. for mech* |  |  |
| *Odocoileus hemionus sitkensis* | y | y |
| *Oncorhynchus* spp*.* | y | y |
| *Oreamnos americanus* | y | y |
| *Peromyscus* spp. | y | y |
| *Phoca vitulina* | y | y |
| *Tamiasciurus hudsonicus* |  | y |
| *Ursus americanus* | y | y |

**SI Figure Legends**

**Figure S1:** Sequence alignment of common, vertebrate prey species for wolves in southeast Alaska. As shown, there are no mismatches between the primers (labeled 12SV5F and 12SV5R) and the target sequences for each species.

**Figure S1**

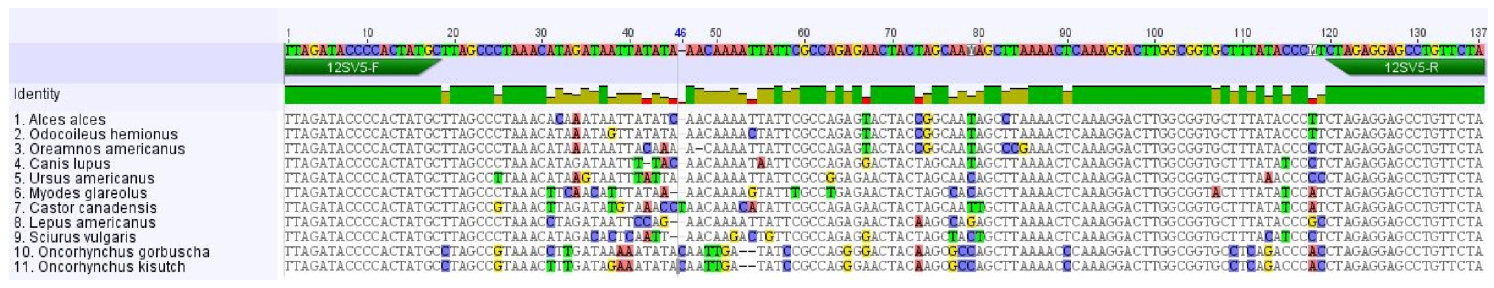
